# North American Powassan virus encompasses diverse *in vitro* phenotypes

**DOI:** 10.1101/2023.08.08.552515

**Authors:** Rebekah J. McMinn, Rose M. Langsjoen, Erica Normandin, Samuel D. Stampfer, Pardis C. Sabeti, Anne Piantadosi, Gregory D. Ebel

## Abstract

Powassan virus (POWV) is a tick-borne flavivirus which has resulted in increasing human cases over the past two decades. Despite high prevalence in ticks and evidence of broad distribution in North America, fewer than 50 human cases are detected annually with evidence of undetected asymptomatic infections. Experimental studies of the relationships between POWV genetic diversity and disease potential are currently lacking. In the present study, sixteen isolates originating from 13 locations in the United States and Canada were used to assess *in vitro* phenotypic diversity in human neuronal cells. Broad differences in replication and cytopathic ability were observed between isolates, even amongst those in the same sublineage. *In vitro* phenotype was not associated with geographic or temporal location and could not be associated with specific genotypes. These results support the observation that the North American POWV population may be highly genetically and phenotypically diverse. The degree to which *in vitro* phenotype reflects transmission and pathogenesis remains to be determined.

## Introduction

Powassan virus (POWV) is an emerging tick-borne flavivirus that causes human neuroinvasive disease in North America. In the United States, reported cases have increased from 0.9 cases per year (1958-2007) to 16.7 cases per year (2008-2021)^1,2^. Though increased recognition and diagnostic capability have undoubtably played a part in this apparent increase, several observations indicate increased virus prevalence: elevated seroprevalence in deer^3^, documented expansion of POWV-associated vectors^4,5^, and reports of high tick infection rates^6,7^. Human cases of POWV largely occur in New England and midwestern states such as Wisconsin and Minnesota. However, the geographic range of POWV is likely significantly broader: POWV has been detected in Colorado, California, Alaska, and New Mexico^8–10^. Moreover, the ecology of POWV and the determinants that lead to local emergence are poorly understood.

POWV is the sole North American member of the tick-borne encephalitis complex (Figs. 1A and C): a group of tick-borne flaviviruses with similar antigenic properties and vector associations^11^. Tick-borne encephalitis virus (TBEV) is broadly distributed throughout Europe and Asia and is divided into genetic subtypes linked to geography, vector species, and pathogenicity in humans. In far-eastern Russia and Asia, TBEV is linked to severe neurological disease resulting in permanent damage to the brain and spinal cord, and death in 20-30% of cases. In Siberia, TBEV causes more mild neurological disease associated with infrequent chronic infection. European TBEV generally causes biphasic febrile disease which can be mildly neurological in the second phase^12^. Case fatality rates for Siberian and European subtypes are between 6-8% and 1-2% respectively. These observations have led us to speculate that similar phenotypic diversity may exist among North American strains of POWV.

**Figure 1.**
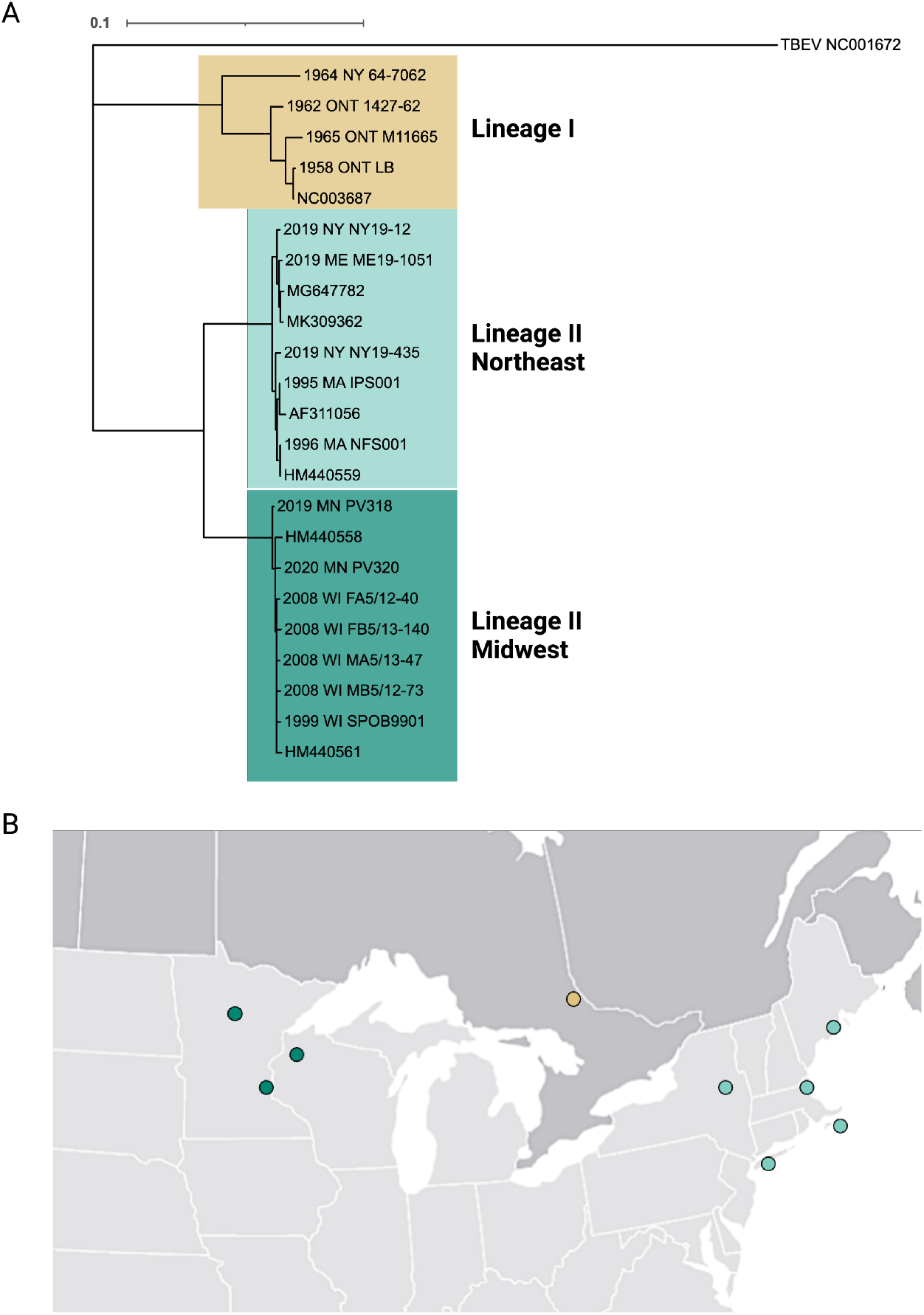
POWV isolates used. A) Maximum likelihood of sixteen POWV and DTV isolates used in this study including several published reference strains and TBEV-EU used as an outgroup. B) Approximate locations of isolates in the U.S. and Canada. Dots may represent multiple isolates in proximity. Created with Biorender.com.

There are two genetic lineages of POWV that have 84.6% nucleotide identity and 95.1% amino acid identity (Figs. 1A and 2) and are indistinguishable by serology. Both virus lineages can cause disease in humans, but lineage II (deer tick virus; DTV) has been detected in humans more frequently due to its association with the aggressively human biting tick *Ixodes scapularis*^13–15^. Clinical disease ranges from asymptomatic infection to severe neurological involvement, with 50% of patients developing permanent disorders and 11% succumbing to disease^1,13,16,17^. Although a range of clinical presentations is associated with POWV infection in people, the severity of both acute clinical disease and the high prevalence of neurological dysfunction among survivors are extremely concerning.

Phenotypic studies of POWV to date have almost exclusively used two highly passaged historical isolates: LB (lineage I) and SPO (lineage II). Therefore, we used 16 low-passage virus isolates collected from human and ticks from Northeast and Midwest U.S. and Canada to assess phenotypic diversity of POWV *in vitro*. Using human neuroblastoma cells, we assessed replication and cytopathic effects to understand how individual POWV strains might differ in their pathogenic potential. We performed full virus genome sequencing pre- and post-passage to identify single nucleotide variants (SNVs) that may be of functional importance. Our results provide clear evidence of significant phenotypic diversity in North American POWV and support the observation that naturally circulating viruses may have distinct propensities for perpetuation, transmission, and pathogenesis.

## Methods

### Cells and viruses

BHK-21 (baby hamster kidney, ATCC CCL-10) cells were maintained in DMEM supplemented with 5% FBS and 1% Penicillin and Streptomycin. SH-SY5Y (human neuroblastoma, ATCC CRL-2286) cells were maintained in a 1:1 ratio of EMEM:DMEM F12 supplemented with 10% FBS and 1% Penicillin and Streptomycin. Cells were grown at 37ºC with 5% CO^2^. Cells were not maintained past passage 50. A table of the POWV isolates used in this study are in Table 1. All POWV stocks were amplified in BHK cells for three to six days, aliquoted with additional 20% FBS and stored at -80ºC. Virus was tittered via plaque assay on BHK cells as described below.

**Table 1.** POWV isolates used in this study. Virus isolates have been obtained either through collaborators or our own tick collections. All viruses have been grown up in BHK cells in our lab to create stocks used in this study. SM = suckling mice, V = Vero cells, B = BHK cells.

| Label | Lineage | Location | Year | Source | Passage History | GenBank accession |
| --- | --- | --- | --- | --- | --- | --- |
| LB | I | Powassan, Ontario | 1958 | human CSF | SM4V2B1 | NC003687 |
| 1427-62 | I | North Bay Ontario | 1962 | *I. marxi* | SM3V1B1 | OP823405 |
| 64-7062 | I | New York | 1964 | *Ixodes sp.* | SM6V2B1 | HM440563 |
| M11665 | I | Laurier Township, Ontario | 1965 | *I. cookei* | SM2V1B1 | OP823404 |
| IPS001 | II | Ipswich, MA | 1995 | *I. scapularis* | SM1B2 | OP823421 |
| NFS9601 | II | Nantucket, MA | 1996 | *I. scapularis* | SM1B2 | HM440559 |
| SPOB9901 | II | Spooner, WI | 1999 | *I. scapularis* | SM1V2B1 | OP823482 |
| FB5/13-140 | II | Spooner, WI site B | 2008 | *I. scapularis* | B2 | OP823484 |
| MA5/13-47 | II | Spooner, WI site A | 2008 | *I. scapularis* | B2 | OP823478 |
| FA5/12-40 | II | Spooner, WI site A | 2008 | *I. scapularis* | B2 | OP823475 |
| MB5/12-73 | II | Spooner, WI site B | 2008 | *I. scapularis* | B2 | OP823480 |
| NY19-12 | II | Saratoga Springs, NY | 2019 | *I. scapularis* | B1 | OP823461 |
| NY19-435 | II | Connetquot, NY | 2019 | *I. scapularis* | B1 | OP823469 |
| ME19-1051 | II | Rockland, ME | 2019 | *I. scapularis* | B1 | OP823442 |
| PV318 | II | St. Croix MN | 2019 | *I. scapularis* | B2 | OL695842 |
| PV320 | II | Northern MN | 2020 | human CSF | B2 | OL695841 |

### Plaque assays

BHK cells were seeded to 90% confluency the day before in six-well cell culture plates. Samples were thawed rapidly and serially diluted in DMEM supplemented with 2% FBS and 1% Penicillin and Streptomycin. All media was removed from six-well plates before BHK cells were inoculated with 200μL of each dilution and incubated at RT on a rocker for one hour. Four milliliters of tragacanth/media overlay was added to each well and incubated at 37ºC for five days. Cells were stained with crystal violet to visualize and count plaques.

*Growth curves*. BHK and SH-SY5Y cells were seeded in six-well cell culture plates the day before for ∼70% confluency. One plate of each cell type was counted the day of to calculate the average cells per well. Virus was diluted to inoculate each cell type at a multiplicity of infection (MOI) of 0.01. Media was removed from each cell type, 200μL inoculum was added to the cells and incubated at RT on a rocker for one hour. Cells were washed three times with PBS and four milliliters of cell media with decreased FBS concentration of 2% was added to each well. The first time point (day 0) was taken immediately before plates are incubated at 37ºC with 5% CO^2^ for the remainder of the experiment. The remainder of the unused virus inoculum was used to immediately perform back titrations via plaque assay. For the remaining five to nine days, supernatant samples were collected from each well in 24-hour increments and stored at -80ºC. Plaque assays were performed as described above to measure infectious virus titer over time. Growth curves were performed in triplicate for both cell types. Growth curves were repeated for SH-SY5Y cells and data was combined for a total of six replicates per isolate. Tests for overall statistical significance were performed in GraphPad Software version 9.5.1 (San Diego, California) for two and six dpi timepoints in SH-SY5Y cells. Kruskal-Wallis test for statistical significance was performed for lineage I and lineage II Midwest. Mann-Whitney t-test was performed for lineage II northeast.

### CPE assay

Cells were seeded to 70% confluency in 96-well cell culture plates. Viral stocks were diluted 1×10^3^ PFU/mL in DMEM supplemented with 2% FBS and 1% Pen/Strep. Diluted virus (50μL) was transferred to the cell culture plates and incubated for up to nine days. For each time point, 20μL of 5mg/mL MTT live cell stain was added to each well and incubated at 37ºC for three hours. The media was removed, and cells were washed twice with PBS. 100μL of 1:2 DMSO:EtOH was added to solubilize the MTT crystals and incubated overnight in the dark at 4ºC. Absorbance was measured at a wavelength of 490nm with background subtraction. Absorbance is normalized to negative control wells on each plate and calculated as a percent cytopathic effect (CPE). Kruskal Wallis test with Dunn’s multiple comparisons was performed in GraphPad Software version 9.5.1 (San Diego, California).

### Sequencing and iSNV analysis

POWV isolates (Table 1) were grown in triplicate on SH-SY5Y cells for nine days. RNA was extracted from 50μL of clarified day nine and input supernatant using the Mag-Bind Viral DNA/RNA kit on a KingFisher Flex Purification System. POWV genome sequencing was performed on RNA as previously described ^18^ Briefly, samples underwent DNase (ArcticZymes), single-cycle cDNA synthesis using random primers and Superscript III (Invitrogen), library tagmentation and 16-cycle PCR amplification using Nextera XT (Illumina), and 150 bp paired-end sequencing using an Illumina MiSeq or NextSeq.

Duplicate independent libraries underwent deep sequencing for analysis of intrasample single nucleotide variants (iSNV). Trimmed, quality-filtered .bam files were generated using viral-ngs v2.0.21, and a merged .bam file was generated using samtools v1.10 (Li et al. 2009). Bam files were separated into respective R1 and R2 fastq files using samtools. Reads <25 bases in length and PCR duplicates were removed using fastp v0.23.2^19^. For each library, fastq files for both tick-derived and BHK-derived RNA were aligned to their respective tick-derived consensus sequence using bowtie2 v2.3.5.1^20^ using the following parameters: --local -L 25 -N 1 --gbar 15 - -rdg 5,1 --rfg 5,1 --score-min G,30,15 --mp 10 (Table S1). Output sam files were converted to sorted, indexed bam files using samtools, and iSNVs were called with VPhaser-2 v2.0 ^21^ with the following parameters: -ps 100 -ig 20 -dt 0 -a 0.001. Allele information for positions present in both libraries was extracted from the corresponding merged bam file. iSNVs were then filtered for those present at ≥ 1% allele frequency and indels were additionally filtered for those greater than two nucleotides in length to reduce spurious iSNV calls. Remaining iSNVs were manually inspected and any suspected spurious calls were removed. iSNVs were considered spurious if: 1) iSNV only occurred in one position across multiple reads (e.g. only at left-most part of reads and never in the middle of reads); 2) iSNV only occurred in one mapping orientation; 3) iSNV only occurred in combinations with one another suggestive of processing artifact. Shannon’s entropy was calculated as follows:

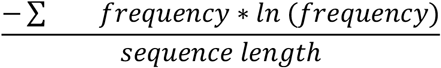

Normal distribution was assessed by Q-Q plot, and significance was assessed by Wilcoxon sign-rank test in SPSS version 28 (IBM Corp., Armonk, New York). Associated amino acid changes were annotated using custom genome annotator excel files, and nucleotide positions were indexed to the HM440559.1 genome. Protein structure depictions of POWV mutations were generated using PyMol (http://pymol.org) and modeled using the cryo-EM structure of TBEV full-length env protein, PDB ID 5O6A. All residues of interest are conserved between POWV and TBEV env (with 78.1% overall identity).

## Results

### POWV isolate characteristics

Sixteen POWV and DTV isolates were chosen as a representative sample set to assess replication phenotypes *in vitro* (Table 1). These included four lineage I and twelve lineage II isolates (five from the Northeast clade, and seven from the Midwest clade). The lineage I isolates (1958-1965) derive from human (12.5%) and tick (87.5%) sources from Ontario, Canada, and New York. The lineage II isolates (1995-2020) originate from five U.S. states and were all isolated from ticks, most of which were *I. scapularis* (78.6%). MN20-CSF01 was isolated from a human patient. Isolates were maintained at the lowest possible passage; however, many of the older isolates have been passaged multiple times through suckling mice.

The 16 isolates used in this study have an overall nucleotide identity of 91.6% (97.0% amino acid identity). Lineage I isolates are the most dissimilar to each other with 94.4% identity, and at least 105 single nucleotide variants (SNVs) between them. Lineage II isolates are overall 96.5% identical: 99.4% among Northeast isolates and 99.6% among Midwest isolates (Table S2). In the Midwest DTV group, SPOB9901 and MA5/13-47 are the most similar with 14 SNVs between them and a single amino acid difference in the NS5 RNA-dependent RNA polymerase (RDRP) at position 8819.

### Virus replication and cytopathic effects in mammalian cells

Initial phenotypic analysis was performed on a subset of the isolates from Table 1. Two mammalian cells lines: baby hamster kidney (BHK-21, ATCC CCL-10), and human neuroblastoma (SH-SY5Y, ATCC CRL-2266) were used to assess replication phenotypes. A multiplicity of infection (MOI) of 0.01 was used to attain multi-step replication. In BHK-21 cells, viral loads in the supernatant peaked at three days post infection (dpi) and universal cell mortality was visually observed three to five dpi.

Replication curves were uniform among lineage II / DTV isolates, though variation was observed between lineage I isolates: LB and 64-7062, the most highly passaged isolates, peaked two dpi around 10^7^ PFU/mL, while M11665 and 1427-62 peak three dpi around 10^6^ PFU/mL (Fig 3A). In SH-SY5Y cells, LB, 64-7062, M11665 replicated similarly, peaking two dpi around 10^7^-10^8^ PFU/mL; however, delayed replication was observed for 1427-62 which peaked six dpi around 10^6^ PFU/mL. DTV replication was highly variable in SH-SY5Y cells with virus titer ranging from 10^2^ to 10^6^ PFU/mL at two dpi. All DTV isolates peaked on six dpi with viral titers ranging from 10^5.5^ to 10^8^ (Fig. 3B).

**Figure 3.**
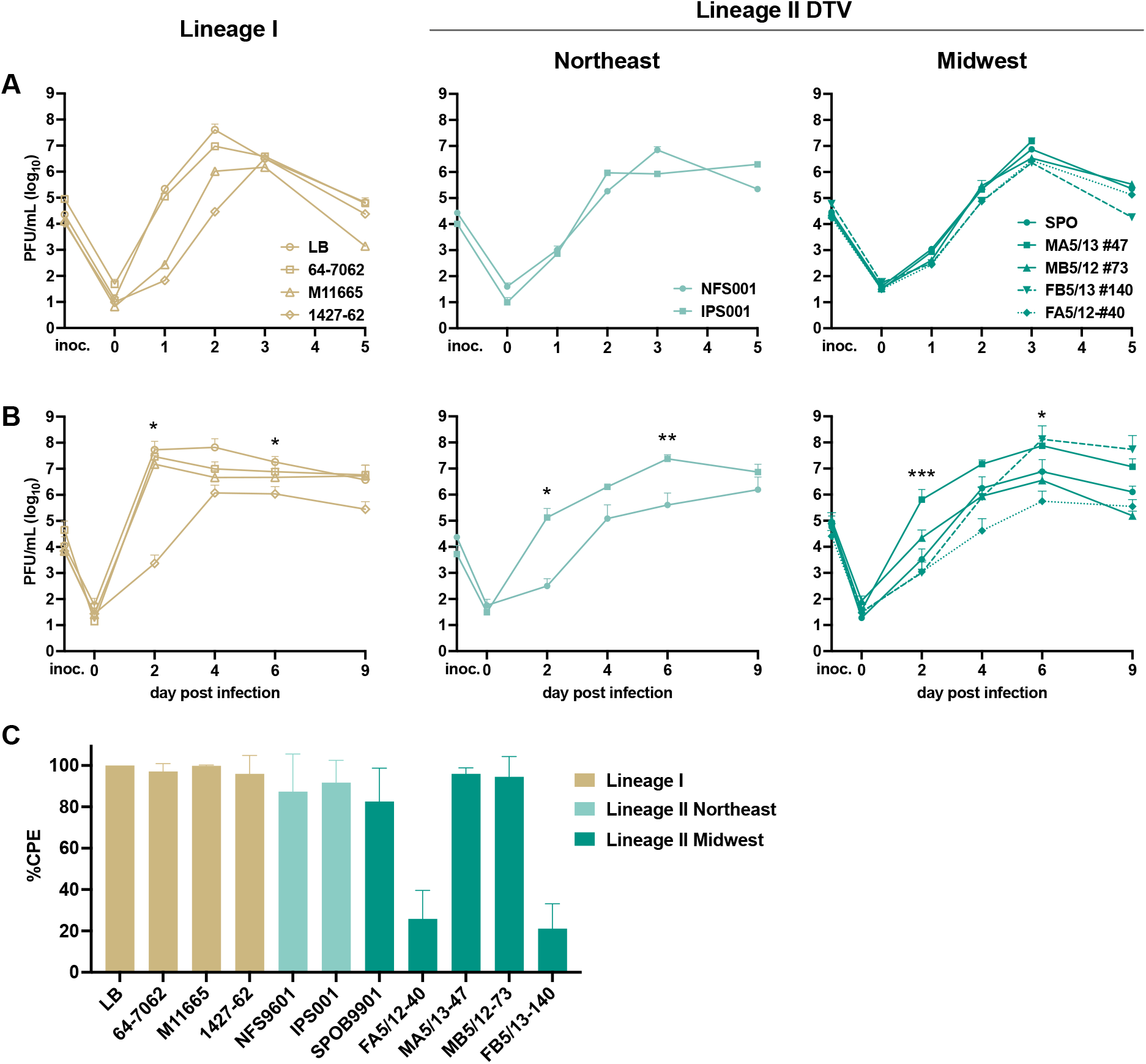
Replication and CPE phenotypes of POWV isolates 1958-2008. POWV is inoculated at an MOI of 0.01 in triplicate on A) BHK and B) SH-SY5Y for incubated up to 9 days at 37ºC. Replication curves are separated by genetic clade to appreciate the differences between closely related isolates. Data shown is from two replicate experiments. C) SH-SY5Y cells are inoculated (n=6 wells) with virus approximating an MOI of 0.01 and incubated for 9 days. Cells are stained with MTT, and absorbance is measured. Absorbances are background subtracted and normalized to uninfected control wells to calculate %CPE. Data shown is from two replicate experiments.

Variable CPE was visually noted during infection in SH-SY5Y cells, thus a cell viability assay was used to measure CPE nine dpi. FA5/12-40 and FB5/13-140 (lineage II Midwest) produced less CPE compared to the other isolates, though significance was only determined against lineage I isolates (p<0.01, Kruskal Wallis test). Lineage I isolates, MA5/13-47, and MB5/12-73, NFS9601, IPS001, and SPOB9901 resulted in 100% CPE (Fig. 3C).

To bolster observations of *in vitro* phenotype, additional current low-passage isolates were obtained (those from 2019-2020), and replication and CPE assays were repeated in SH-SY5Y cells. Though the shape of the replication curves was different, significant variability was again observed between isolates. Highly variable replication was observed with peak viral titer ranging from 10^4.5^ to 10^8.5^ PFU/mL. Isolates from New York (NY19-435 and NY19-12) resulted in much lower titers than the isolates from Minnesota (MN19-61 and MN20-CSF01) and Maine (ME19-1051) (Fig 4A). All these isolates caused 100% CPE by nine dpi (Fig 4B).

**Figure 4.**
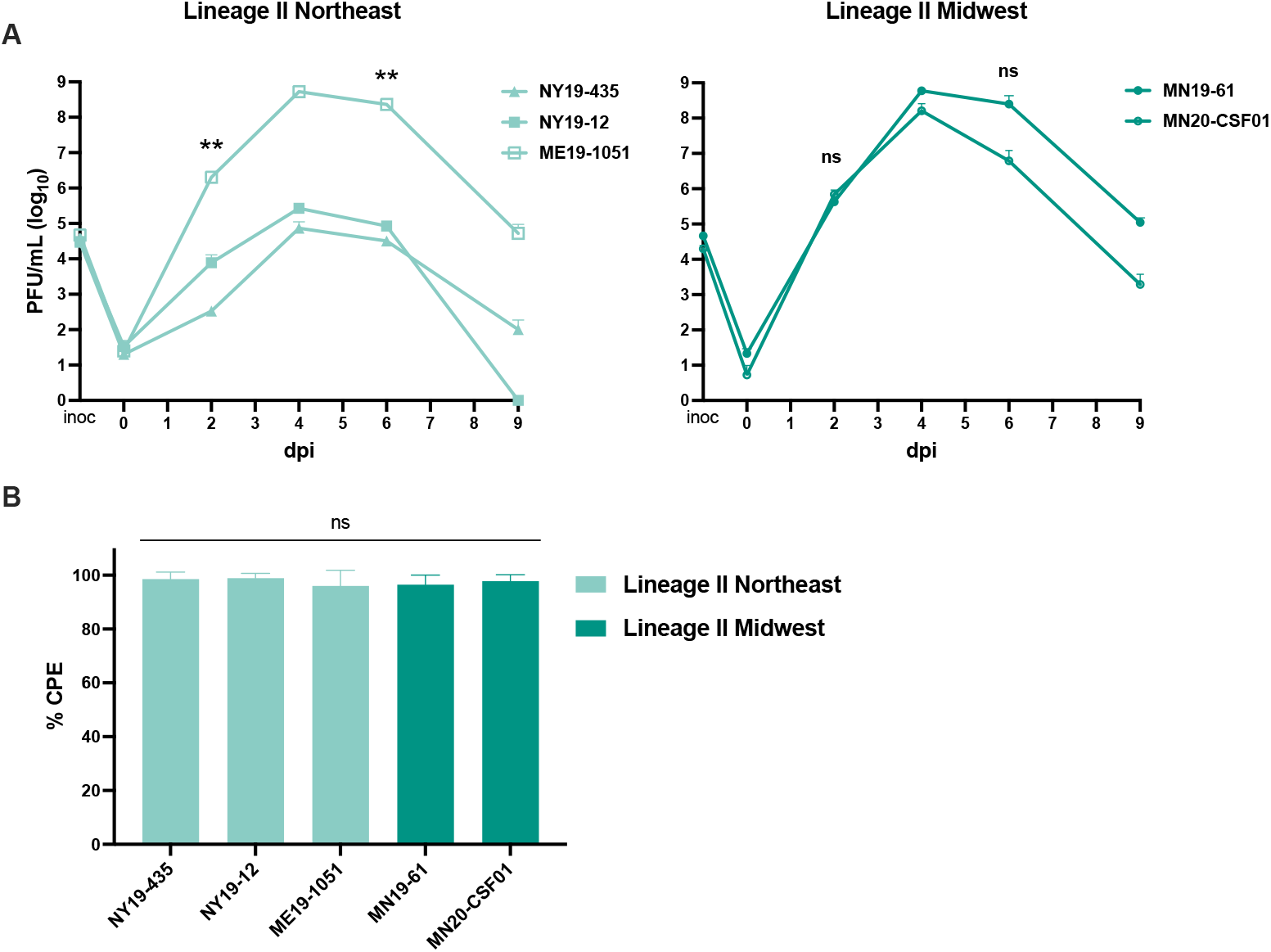
A) Replication and B) CPE of 2019-2020 DTV isolates on SH-SY5Y cells. Results are from a single experiment.

### Potential genetic determinants of phenotype

To assess potential genetic links to phenotype, we identified consensus variants shared between multiple DTV isolates of the same genetic clade. No variant was shared by more than two isolates and could not be linked to *in vitro* phenotype.

Full viral genome sequencing was performed on RNA isolated from pre-infection ‘input’ POWV and after nine days replication in SH-SY5Y cells to identify mutations that may be associated with *in vitro* phenotype and replication in neuronal cells. Isolates included in this experiment were LB, 64-7062, M11665, 1427-62, NFS9601, IPS001, MA5/13-47, MB5/12-73, and FB5/13-140. Eleven total consensus-level mutations occurred after a single passage in SH-SY5Y cells, ranging from zero to five mutations per isolate (Fig 5A). Seven of these eleven mutations occurred in the env protein, all of which were nonsynonymous. Two nonsynonymous mutations in env occurred in multiple isolates: G1096A was identified in lineage I isolates 1427-62 and M11665, and A1868G was identified in lineage II isolates MA5/12 #47, SPOB9901, and IPS001. In most cases, the env mutation was not found as even a minor variant in the input sample, and rose to high frequency by day 9, suggesting possible selection (Fig. 5A).

**Figure 5.**
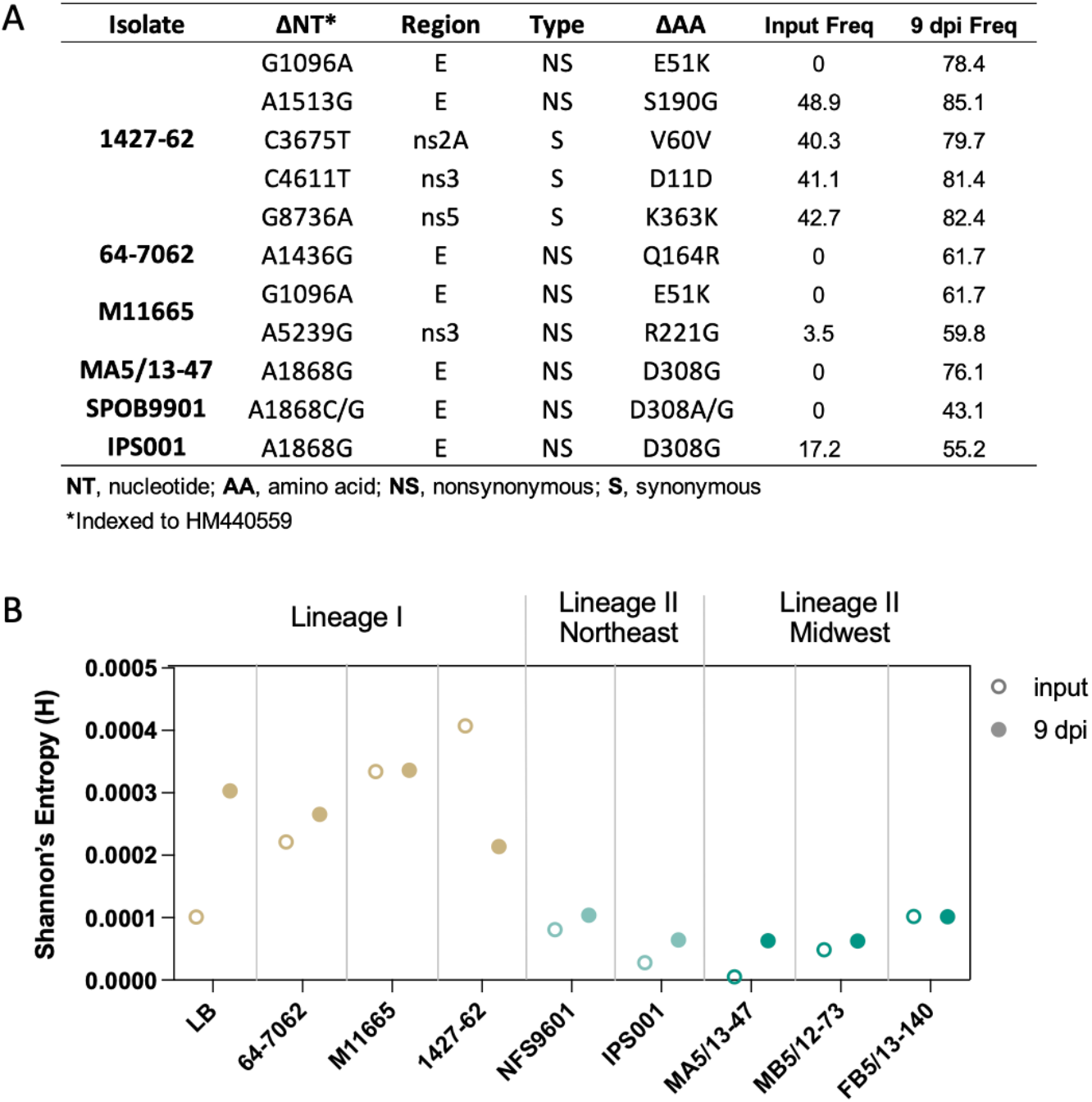
Intra-isolate genetic changes pre- and post-SH-SY5Y culture. A) All consensus level SNVs identified after a single passage in SH-SY5Y cells. B) Intra-isolate complexity pre- and post-SH-SY5Y culture, as measured by Shannon’s entropy from merged biological replicate libraries. (Not significant by Wilcoxon signed rank test, two-tailed, p=0.13

We next evaluated intra-sample single nucleotide variants (iSNVs) to describe changes in quasispecies diversity between input and nine dpi SH-SY5Y culture samples. All lineage I isolates started with greater diversity as measured by average Shannon’s entropy than lineage II isolates, perhaps due to higher passage history. All isolates except for 1427-62 increased in diversity over 9 days of passage in SH-SY5Y cells, though these changes were not statistically significant (Fig. 5B).

## Discussion

POWV has emerged in recent decades due to the expansion of deer ticks and increased recognition and diagnosis of disease in humans. However, the incidence of human infection, while increasing, lags far behind other areas where similar viruses (e.g. TBEV) have emerged.

The reasons for this are unclear at present but is of significant interest due to the clinical severity of POWV disease in human beings and the ecological changes to the North American landscape that have facilitated the emergence of other tick-borne pathogens. In this work we sought to assess the overall hypothesis that the relative paucity of human cases may be due to reduced pathogenic potential of some currently circulating isolates. Specifically, we measured *in vitro* replication, evaluated CPE in neuronal cells, and assessed virus genetics during replication in cultured cells.

Sixteen viral isolates from three genetic clades were used in the present study. The represented lineage I isolates are more genetically dissimilar than the DTV isolates. DTV genetic subclades are tied to geographic location: Northeast and Midwest U.S. Many of the lineage I and II isolates in current use are moderately to highly passaged; therefore, several current low-passage isolates from a broad geographic range were included in this study. The five DTV isolates from Spooner, WI present a unique opportunity to assess phenotype between isolates with close geographic and temporal characteristics. While SPO9901 was collected in 1999 and has been moderately passaged, it has similar consensus identity to the four low-passage isolates collected in 2008 (99.8%), indicating genetic stability of DTV in this area as previously described^22^. Despite the geographic proximity, they are genetically distinct, with 14-33 consensus differences among them. DTV isolates from 2019 originate from *I. scapularis* collected from four locations and represent both the Northeast and Midwest (Table 1 and Fig. 1C). Out of the 16 isolates evaluated in this study, two isolates originate from cerebral spinal fluid of patients with POWV disease. Together, these isolates represent a broad geographic and temporal range from all genetic clades, many of which are maintained with few passages in cell culture. Thus, the isolates included in this study are suitable to assess *in vitro* phenotypes.

SH-SY5Y cells are frequently used as *in vitro* disease model for encephalitic flaviviruses as they are neuronal cells originated from a bone-marrow-derived neuroblastoma. Here, we quantify virus replication and cytopathic effects in SH-SY5Y cells as a proxy for pathogenic potential in humans. BHK-21 cells are kidney-derived fibroblasts used extensively as general laboratory cells, highly permissive to flavivirus replication, and used here as a robust comparison of virus replication.

Isolates from all genetic clades were able to replicate to high titers in bone marrow-derived SH-SY5Y neuroblastoma cells, though significant variation was observed. At two dpi, over a 10,000-fold (lineage I, lineage II Northeast) and 1000-fold (lineage II Midwest) difference in viral titer was observed. Among DTV isolates, 10,000-fold differences were observed in peak viral titers at four or six dpi (Figs. 3B and 4A). Surprisingly, no correlation between replication phenotype and genetic clade, location, or originating source were observed. These observations clearly establish that POWV isolates possess extreme variation in their ability to replicate in neuronal cells.

Inspection of cell mortality nine dpi in human neuronal cells revealed virus strain-dependent differences in CPE. Most isolates resulted in 100% CPE on SH-SY5Y cells, apart from FB5/13-140 and FA5/12-40 (lineage II Midwest; Fig 3C). Interestingly, FB5/13-140 and FA512-40 had both the highest and lowest peak viral titers (respectively) of the DTV Midwest isolates, indicating CPE phenotype did not correlate with increased replication (Fig. 3B). To determine whether any recent DTV isolates would be similarly unable to cause CPE, we repeated this experiment with five DTV isolates from 2019-2020. These isolates caused CPE in SH-SY5Y cells quite efficiently, suggesting that rarely, POWV strains cause minimal CPE in these cells. Collectively, these experiments document strain-dependent variation in the ability of POWV to cause CPE in a neuronal cell *in vitro* model. The significance of these data for human disease remains unclear at present.

Given the high passage history of some of the isolates used, enhanced replication could be due to adaptations to cell culture or suckling mouse brain. Inoculation into the brains of suckling mice was (and sometimes still is) used to isolate POWV, though currently cell culture is used more frequently. For instance, lineage I isolates have been passaged multiple times in suckling mice and nearly all (excluding 1427-62) replicate similarly well in SH-SY5Y cells, despite having only 95.6% nucleotide identity. Thus, efficient replication could be due to adaption to neuronal cells. In contrast, the following observations demonstrate DTV phenotypes in SH-SY5Y cells are not a result of adaptation to cell culture: (1) uniform replication and CPE of DTV isolates in BHK cells, despite some being more highly passaged, and (2) high variation in SH-SY5Y cells between isolates with similar passage history. Most of the studies on POWV to date have used isolates LB (highly passaged) and SPO (moderately passaged) in experimental studies on transmission and disease. The present results demonstrate the importance of working with low-passage isolates for an accurate picture of POWV phenotypes.

Two nonsynonymous consensus changes were identified in post-SH-SY5Y culture samples that are shared between multiple isolates: G1096A (*env* E51K) and A1868G (*env* D308G). Structural analysis of *env* residues E51 and D308 were performed using the cryo-EM structure of TBEV *env*. Interestingly, D308 lies on the outer surface of domain III in the protein binding motif (Fig. 6A) and has been implicated for increased neuropathic potential in TBEV^23^. Residue E51 lies in domain I of the env protein, which is involved in protein dimerization and membrane fusion^23^, though this residue has not been associated with neurovirulence. The equivalent region (residues 51-54) is characterized as the hinge region in Dengue given its proximity to domain II (which contains the fusion loop), and mutations in the region can result in fusion being triggered at a higher pH^24^. E51 is positioned alongside K280 in TBEV env (Fig. 6B) and may form a stabilizing hydrogen bond, which the E51K mutation would change to a repulsion to push the two domains apart, possibly destabilizing the prefusion env conformation to allow membrane fusion to occur more easily. This region contains nearby histidines 275 and 282 that appear to be part of the histidine switch mechanism which triggers membrane fusion, wherein protonation of adjacent histidines at low pH results in new charge-charge repulsions that induce conformational changes to mediate membrane fusion^25^. This suggests that the dominance of E51K (G1096A) *in vitro* may be due to having fusion more easily triggered through destabilization of the prefusion conformation of env, a feature that may not be as favorable *in vivo. In vitro* phenotype could not be linked to either SNV, thus additional studies are needed to determine the significance of these results.

**Figure 6.**
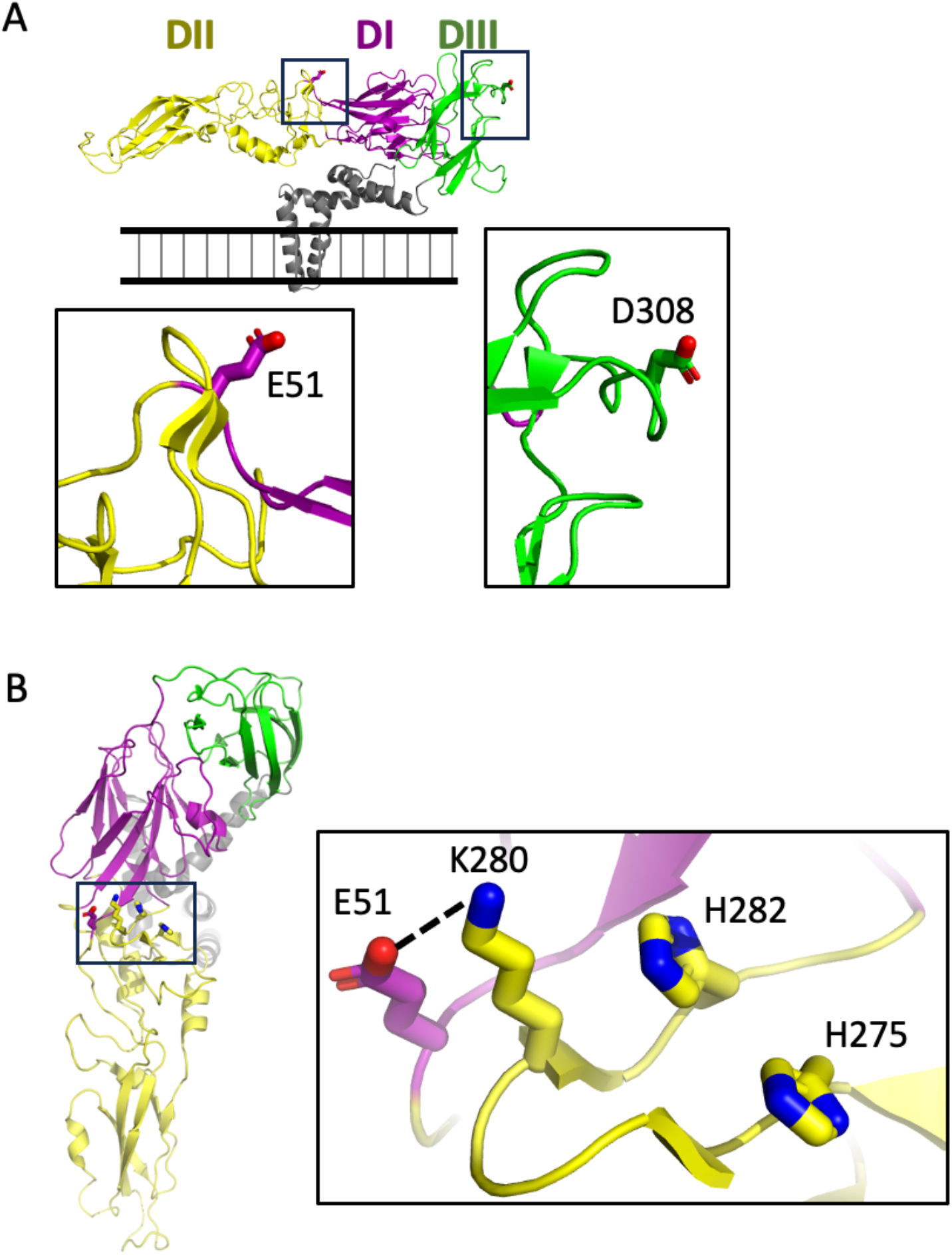
E51K and D308G mutations. A) E51 and D308 as seen on env protein from TBEV, side view of the molecule in the membrane. B) Possible intramolecular hydrogen bond between E51 and K280 would change to a repulsion with E51K mutation, creating additional repulsions in the setting of low pH with positively-charged histidines 275 and 282, as seen from top view looking down at the exterior of a virion. Modeled on TBEV whole virion cryo-EM structure PDB ID 5O6A^26^ all sidechains shown are identical in Powassan virus env.

In summary, our results demonstrate extensive *in vitro* phenotypic diversity present in POWV. Further investigation using *in vivo* models is needed to determine how these results may translate to disease in humans. Though no clear genetic determinants of *in vitro* phenotype were observed, there are a few key residues that warrant further investigation. Thus, the replication of POWV in human neuroblastoma cells and the ability of strains to cause CPE is highly variable and not currently attributable to a key set of viral genetic determinants.

## References

1. Ebel, G. D. Update on Powassan Virus: Emergence of a North American Tick-Borne Flavivirus. Annu. Rev. Entomol. 55, 95–110 (2010).

2. Powassan virus. (2022).

3. Nofchissey, R. A. et al. Seroprevalence of Powassan virus in New England deer, 1979-2010. Am. J. Trop. Med. Hyg. 88, 1159–1162 (2013).

4. Eisen, R. J., Eisen, L. & Beard, C. B. County-Scale Distribution of Ixodes scapularis and Ixodes pacificus (Acari: Ixodidae) in the Continental United States. doi:10.1093/jme/tjv237

5. Dennis, D. T., Nekomoto, T. S., Victor, J. C., Paul, W. S. & Piesman, J. Reported Distribution of Ixodes scapularis and Ixodes pacificus (Acari: Ixodidae) in the United States. J. Med. Entomol. 35, 629–638 (1998).

6. Klinepeter, K. Powassan Virus Identified in Ticks. (2022).

7. Campbell, O. & Krause, P. J. The emergence of human Powassan virus infection in North America. Ticks Tick. Borne. Dis. 11, 101540 (2020).

8. Deardorff, E. R. et al. Powassan Virus in and New Mexico, USA, and Russia, 2004–2007. Emerg. Infect. Dis. 19, 1–5 (2013).

9. Johnson, H. N. Isolation of Powassan virus from a spotted skunk in California. J. Wildl. Dis. 23, 152–153 (1987).

10. Thomas, L. A., Kennedy, R. C. & Eklund, C. M. Isolation of a Virus Closely Related to Powassan Virus from Dermacentor andersoni Collected along North Cache la Poudre River, Colo. https://doi.org/10.3181/00379727-104-25836 104, 355–359 (1960).

11. Calisher, C. H. et al. Antigenic relationships between flaviviruses as determined by cross-neutralization tests with polyclonal antisera. J. Gen. Virol. 70, 37–43 (1989).

12. Gritsun, T.., Lashkevich, V.. & Gould, E.. Tick-borne encephalitis. Antiviral Res. 57, 129–146 (2003).

13. Piantadosi, A. et al. Emerging Cases of Powassan Virus Encephalitis in New England: Clinical Presentation, Imaging, and Review of the Literature. Clin. Infect. Dis. 62, 707–713 (2016).

14. Normandin, E. et al. Powassan Virus Neuropathology and Genomic Diversity in Patients With Fatal Encephalitis. Open Forum Infect. Dis. 7, (2020).

15. El Khoury, M. Y. et al. Potential Role of Deer Tick Virus in Powassan Encephalitis Cases in Lyme Disease-endemic Areas of New York, USA. Emerg. Infect. Dis. • http://www.cdc.gov/eid • 19, (2013).

16. Krow-Lucal, E. R., Lindsey, N. P., Fischer, M. & Hills, S. L. Powassan Virus Disease in the United States, 2006–2016. Vector-Borne Zoonotic Dis. 18, 286–290 (2018).

17. R, D., S, S. & JP., W. Arboviruses in New York State: an attempt to determine the role of arboviruses in patients with viral encephalitis and meningitis. Am. J. Trop. Med. Hyg. 28, 577–82 (1979).

18. Normandin, E. et al. Powassan virus neuropathology and genomic diversity in patients with fatal encephalitis. Open Forum Infect. Dis. (2020). doi:10.1093/ofid/ofaa392

19. Chen, S., Zhou, Y., Chen, Y. & Gu, J. fastp: an ultra-fast all-in-one FASTQ preprocessor. Bioinformatics 34, i884–i890 (2018).

20. Langmead, B. & Salzberg, S. L. Fast gapped-read alignment with Bowtie 2. Nat. Methods 9, 357–359 (2012).

21. Yang, X., Charlebois, P., Macalalad, A., Henn, M. R. & Zody, M. C. V-Phaser 2: Variant inference for viral populations. BMC Genomics 14, 1–10 (2013).

22. Brackney, D. E., Nofchissey, R. A., Fitzpatrick, K. A., Brown, I. K. & Ebel, G. D. Short report: Stable prevalence of powassan virus in Ixodes scapularis in a Northern Wisconsin focus. Am. J. Trop. Med. Hyg. 79, 971–973 (2008).

23. Kellman, E. M., Offerdahl, D. K., Melik, W. & Bloom, M. E. Viral Determinants of Virulence in Tick-Borne Flaviviruses. doi:10.3390/v10060329

24. Butrapet, S. et al. Amino acid changes within the E protein hinge region that affect dengue virus type 2 infectivity and fusion. Virology 413, 118–127 (2011).

25. Fritz, R., Stiasny, K. & Heinz, F. X. Identification of specific histidines as pH sensors in flavivirus membrane fusion. J. Cell Biol. 183, 353–361 (2008).

26. Füzik T, Formanová P, Růžek D, Yoshii K, Niedrig M, Plevka P. Structure of tick-borne encephalitis virus and its neutralization by a monoclonal antibody. Nat Commun. 2018;9(1):1-11. doi:10.1038/s41467-018-02882-0

